# Mixed Model Association with Family-Biased Case-Control Ascertainment

**DOI:** 10.1101/046995

**Authors:** Tristan Hayeck, Noah A. Zaitlen, Po-Ru Loh, Samuela Pollack, Alexander Gusev, Nick Patterson, Alkes L. Price

## Abstract

Mixed models have become the tool of choice for genetic association studies; however, standard mixed model methods may be poorly calibrated or underpowered under family sampling bias and/or case-control ascertainment. Previously, we introduced a liability threshold based mixed model association statistic (LTMLM) to address case-control ascertainment in unrelated samples. Here, we consider family-biased case-control ascertainment, where cases and controls are ascertained non-randomly with respect to family relatedness. Previous work has shown that this type of ascertainment can severely bias heritability estimates; we show here that it also impacts mixed model association statistics. We introduce a family-based association statistic (LT-Fam) that is robust to this problem. Similar to LTMLM, LT-Fam is computed from posterior mean liabilities (PML) under a liability threshold model; however, LT-Fam uses published narrow-sense heritability estimates to avoid the problem of biased heritability estimation, enabling correct calibration. In simulations with family-biased case-control ascertainment, LT-Fam was correctly calibrated (average *χ*^2^ = 1.00), whereas Armitage Trend Test (ATT) and standard mixed model association (MLM) were mis-calibrated (e.g. average *χ*^2^ = 0.50-0.67 for MLM). LT-Fam also attained higher power in these simulations, with increases of up to 8% vs. ATT and 3% vs. MLM after correcting for mis-calibration. In 1,269 type 2 diabetes cases and 5,819 controls from the CARe cohort, downsampled to induce family-biased ascertainment, LT-Fam was correctly calibrated whereas ATT and MLM were again mis-calibrated (e.g. average *χ*^2^ = 0.60-0.82 for MLM). Our results highlight the importance of modeling family sampling bias in case-control data sets with related samples.

## Introduction

Mixed models have become the tool of choice for genetic association studies^1^-^4^, and the challenges caused by case-control ascertainment in studies of unrelated individuals have been understood and addressed^5^-^7^. In addition, the advantages of mixed model association in studies with related individuals are widely recognized^8^. However, none of those studies considered the consequences of family-biased case-control ascertainment, in which cases and controls are ascertained non-randomly with respect to family relatedness. Previous work has shown that family-biased ascertainment can severely bias heritability estimates^9^^;^ ^10^, but the consequences for mixed model association have not previously been investigated. We show that family-biased case-control ascertainment leads to severe biases in mixed model association statistics, and propose a new liability threshold mixed model association statistic for family-based case-control studies (LT-Fam) that is robust to this problem.

In our previous work^5^, we introduced a liability threshold based mixed model association statistic (LTMLM) that addresses the power loss of standard mixed model methods under case-control ascertainment in unrelated individuals. Similar to LTMLM, LT-Fam is computed from posterior mean liabilities (PML) under a liability threshold model conditional on every individual_s case-control status and the disease prevalence. However, LTMLM is susceptible to mis-calibration under family-biased ascertainment, due to biased narrow-sense heritability estimation and calibration based on phenotypic covariance. The LT-Fam statistic is constructed to specifically address family-biased ascertainment, using published narrow-sense heritability estimates and properly controlling for relatedness.

We compared the LT-Fam statistic to ATT and MLM in different settings of family-biased ascertainment by simulating sib pairs under different levels of discordant and concordant sampling. LT-Fam attains proper calibration in simulations with family-biased case-control ascertainment (average *χ*^2^ = 1.00), whereas Armitage Trend Test (ATT) and standard mixed model association (MLM) are both severely mis-calibrated (e.g. average *χ*2 = 0.50-0.67 for MLM). In simulations, LT-Fam attains an increase in power versus existing methods, with increases of up to 8% vs. ATT and 3% vs. MLM after correcting the other statistics for mis-calibration. In 1,269 type 2 diabetes cases and 5,819 controls from the Candidate Gene Association Resource (CARe) cohort, downsampled to induce family-biased case-control ascertainment, LT-Fam is correctly calibrated whereas ATT and MLM are both severely mis-calibrated (e.g. average *χ*^2^ = 0.60-0.82 for MLM).

## Materials and Methods

### Overview of Method

The LT-Fam method consists of three main steps. First, a genetic relationship matrix (GRM) is calculated and then restricted to include only relationships between related individuals by changing GRM entries below a threshold to 0. The narrow-sense heritability is either assumed to be known, or can be estimated in settings without family-biased ascertainment (see Estimation of Narrow-sense Heritability). Second, Posterior Mean Liabilities (PML) are estimated using a truncated multivariate Gibbs sampler. The PMLs are conditional on the relatedness, case-control status of all individuals, and prevalence of the disease (see Posterior Mean Liabilities). Third, a *χ*^2^ (1 d.o.f) association score statistic is computed based on the association between the candidate SNP and the PML. The LT-Fam statistic is constructed using published narrow-sense heritability estimates as well as genetic relatedness using a threshold, as opposed to LTMLM which uses SNP-heritability estimates and calibration based on phenotypic covariance without thresholding (see LT-Fam Association Statistic). We have released open-source software implementing the LT-Fam statistic (see Web Resources).

To better understand the need to account for family-biased ascertainment it is helpful to consider a toy example. Figure 1 depicts (A) the conditional probabilities of being a case given that an individual_s sibling is a case and (B) the probability of being a case given that an individual_s sibling is a control, assuming a 100% heritable trait under a liability threshold model at different values of disease prevalence. Thus, the conditional probability of being a case or a control can be heavily influenced by the disease status of an individual_s relative(s), depending on disease prevalence.

**Figure 1.**
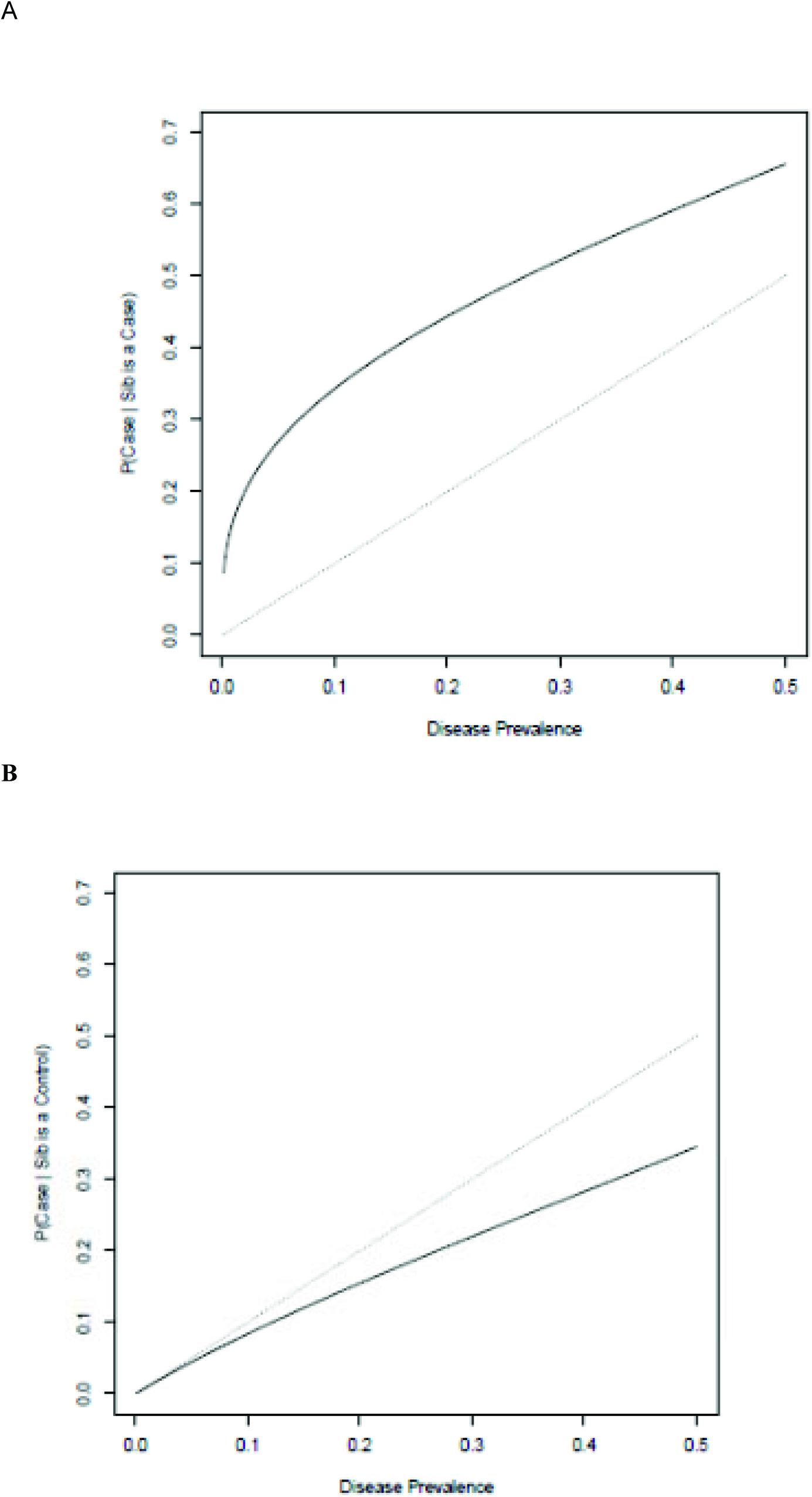
Risk of disease varies with disease status of relatives. Based on analytic derivation of siblings (assuming relatedness of 0.5 and underlying bivariate normal liability), we report (A) the conditional probabilities of being a case given that individual_s sibling is a case and (B) the probability of being a case given the individual_s sibling is a control. The dotted line is the disease prevalence plotted against itself.

### Estimation of Narrow-sense Heritability

Estimating an appropriate heritability parameter is an important step in mixed model association analysis. In studies of related individuals the appropriate heritability parameter is the heritability explained by all genetic variants under an additive model (narrow-sense heritability) *h*^2^.^8^^;^ ^9^^;^ ^11^ In studies of unrelated individuals the appropriate heritability parameter is the heritability explained by genotyped SNPs under an additive model (SNP-heritability) *h*_*g*_^2^ (ref.^3^-^5^), which is generally smaller than *h*^2^. In studies with cryptic relatedness, the appropriate heritability parameter (called “pseudo-heritability” by ref. ^1^) may lie in between *h*_*g*_^2^ and *h*^2^. Since the current work focuses on related individuals, the appropriate heritability parameter is the narrow-sense heritability *h*^2^.

Standard mixed model association methods generally build a genetic relationship matrix (GRM) from genotype data and then estimate a heritability parameter from the phenotypes using restricted maximum likelihood (REML) ^1^-^3^. (The GRM may be constructed excluding the candidate SNP to avoid “proximal contamination_.^3^^;^ ^12^) In studies of related individuals, the GRM can either be constructed from known pedigrees^8^^;^ ^11^ (if available) or directly from the genotype-based GRM by changing GRM entries below a threshold to 0 (thresholded GRM; ref^9^). However, the resulting estimates of *h*^2^ are known to be biased under family-biased ascertainment^9^. Thus, in data sets with family-biased ascertainment, we instead use published estimates of *h*^2^. In this case the LT-Fam statistic will still make use of the thresholded GRM, as described below.

The goal of this work is to test for association between a candidate SNP and a phenotype while controlling for family-biased ascertainment. We first consider a quantitative trait:

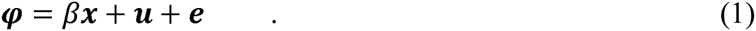

The phenotypic data (transformed to have mean 0 and variance 1) may be represented as a vector *φ* with values for each individual *i*. Genotype values of candidate SNP are transformed to a vector ***x*** with mean 0 and variance 1, with effect size *β*. The quantitative trait value depends on the fixed effect of the candidate SNP (***βx***), the genetic random effect excluding the candidate SNP (***u***), and the environmental component (***e***). We extend to case-control traits via the liability threshold model in which each individual has an underlying, unobserved normally distributed trait called the liability^13^. An individual is a disease case if the liability exceeds a specified threshold *t*, corresponding to disease prevalence and a control if the individual has liability below *t*.

We construct a thresholded GRM^9^

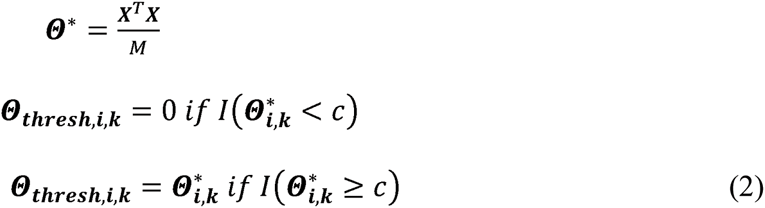

where ***X*** is a matrix of SNPs normalized to mean 0 and variance 1 and *M* is the number of SNPs. We use a threshold of *c*=0.05, as in our previous work^9^.

The phenotypic covariance between individuals is modeled as

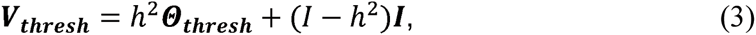

where ***I*** is the identity matrix. As noted above, we estimate *h*^2^ from the data—e.g. via Haseman-Elston (H-E) regression^14^ (or PCGC regression, a more general formulation^15^) followed by transformation to liability scale (Equation 23 of ref ^16^)—if there is no family-biased ascertainment, and we use published estimates of *h*^2^ if there is family-biased ascertainment.

### Posterior Mean Liabilities

The procedure for estimating PML is similar to our published LTMLM method^5^, although the underlying GRM and *h*^2^ parameter are different (see Estimation of Narrow-sense Heritability), as is the way in which the PML is used to compute an association statistic (see LT-Fam Association Statistic).

We first consider the univariate PML (PML_uni_), constructed independently for each individual; we generalize to the multivariate setting below. As described in equations 11 and 12 of ref^16^, these correspond to the expected value of the liability conditional on the case-control status:

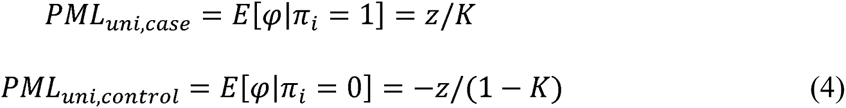

These values are calculated analytically in the univariate setting, and can be thought of as the mean of a truncated normal above or below the liability threshold *t* depending on case-control status^16^.

We now generalize to the multi-variate case:

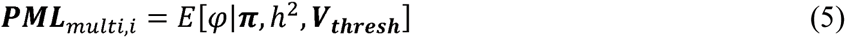

The **PML**_**multi**_ for each individual is conditional on that individual_s case-control status, every other individual_s case-control status, and on the matrix ***V***_***thresh***_. We estimate the PML using a Gibbs sampler, sampling each individual_s liability conditional on the remaining parameters from a truncated multivariate normal distribution. We use 100 burn-in iterations followed by 1,000 additional MCMC iterations, and estimate the PML*_multi_* by averaging over MCMC iterations via Rao-Blackwellization. A summary of the Gibbs sample algorithm is provided below (further details are provided in the LTMLM manuscript^5^):

Initialization: for each individual *i*,

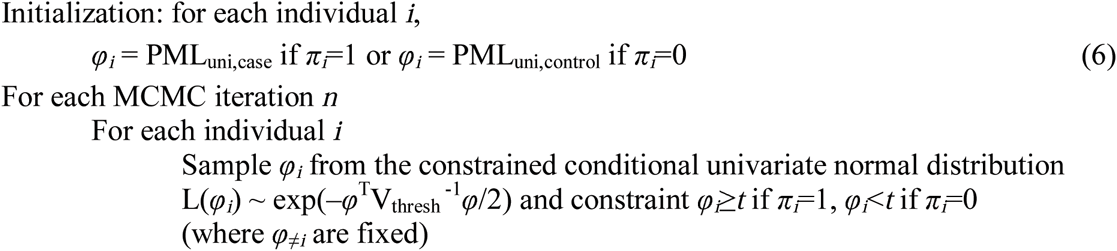

### LT-Fam Association Statistic

The LT-Fam association statistic is a modification of the LTMLM statistic, instead using narrow-sense heritability estimates and ***Θ*_*thresh*_** to control for family-biased ascertainment. The method uses a retrospective association score statistic assuming a liability threshold model. For simplicity, we first consider the case where the liability is known.

We jointly model the liability and the genotypes using a retrospective model, enabling appropriate treatment of sample ascertainment. The score statistic of the joint retrospective model is then (see ref.^5^ for a detailed derivation):

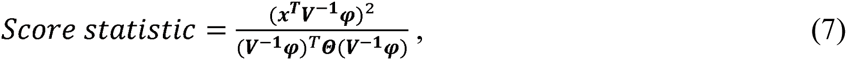

where (in the denominator) ***Θ***, the true underlying genetic relatedness of the individuals, can be approximated by the identity matrix in data sets of unrelated individuals. In the liability threshold setting the liability is approximated using the ***PML*** (analogous to ref.^5^):

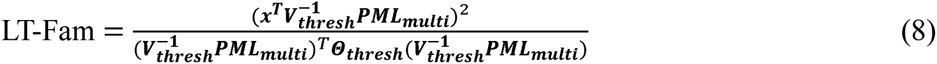

In comparison the ATT, MLM, LTMLM statistics are formulated as:

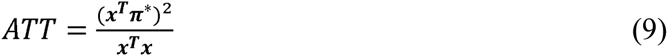

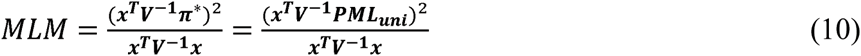

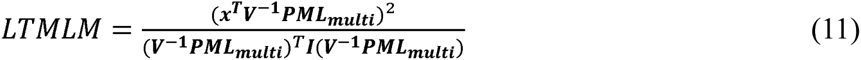

where *π*\* denotes case-control phenotypes normalized to mean zero and variance 1. We note that the LT-Fam and LTMLM use different GRMs (***Θ***_thresh_ vs. ***Θ***), heritability parameters (*h*^2^ vs. *h*_*g*_^2^), and phenotypic covariance matrices ( ***V***_*thresh*_ vs. ***V***). In addition, the denominator of LT-Fam uses the GRM ***Θ***_*thresh*_, as opposed to LTMLM which uses the identity matrix ***I***.

### Simulated Genotypes and Simulated Phenotypes

We performed simulations using simulated genotypes and simulated phenotypes, all involving *N/2* sibling pairs. Three different sibling ascertainment schemes were considered: case-control ascertainment without family bias (unbiased), all concordant siblings, and all discordant siblings. Under each simulation scenario approximately 50% cases and 50% controls were ascertained and 100 separate simulations were run. In runs with *N* = 5,000 a random set of 100 SNPs were set to be causal, and for *N* = 1,000 a random set of 20 SNPs were set to be causal. All simulations included *M* candidate SNPs *(M* = 50,000 or 10,000) and an independent set of *M* SNPs used for calculating the GRM (so that candidate SNPs were not included in the GRM). Half of the causal SNPs were candidate SNPs and the other half were GRM SNPs. Siblings were simulated by generating genotypes of parents for each sib pair, and 25 blocks of SNPs from each parent haplotype were randomly passed along to the children to simulate mating.

Case-control ascertainment (50% cases and 50% controls) was performed using ascertainment probabilities based on the disease prevalence *ƒ*, as follows. Under the unbiased scheme all case-case siblings were retained, case-control siblings were retained with probability *ƒ*\*(1-*ƒ*), and control-control siblings were retained with probability [*ƒ*\*(1-*ƒ*)]. For the concordant scheme, *N/4* sibling pairs were case-case and *N/4* sibling pairs were control-control. For the discordant scheme, all *N/2* sibling pairs were case-control.

The true value of *h*^2^ was set to 0.50 in all simulations. The LT-Fam statistic assumes this parameter to be known (except in the unbiased simulation, in which the H-E-regression estimate is used^14^^;^ ^15^). However, we also performed simulations in which *h*^2^ is incorrectly specified to LT-Fam.

We also performed simulations with shared environment. The environmental term is sampled from a bivariate distribution:

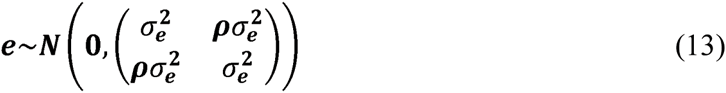

The correlation between siblings was set to *ρ*= 0.75, and the environmental variance was set to 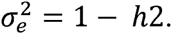.

We compared Armitage Trend Test (ATT), MLM, LTMLM, and LT-Fam statistics. We evaluated performance using average *χ*^2^ statistics at causal, null, and all markers. The AGC at all markers is also resported (median *χ*^2^ divided by 0.455).^17^

### CARe Genotypes and T2D Phenotypes

We analyzed African-American CARe samples with case-control phenotypes for type 2 diabetes (T2D), a disease with prevalence roughly 8%. The data set contained 1,269 cases and 5,819 controls genotyped at 736,614 SNPs after QC^18^. We compared ATT, MLM, and LT-Fam statistics under various downsampling schemes corresponding to unbiased, concordant relative, and discordant relative ascertainment.

First, we analyzed all samples with T2D phenotypes available (unbiased ascertainment). Second, we considered 3 concordant relative schemes: (i) remove cases who do not have a case relative (relatedness > 0.05) in the data set, (ii) remove controls who have a case relative, and (iii) remove both of the previous sets. Third, we considered 3 discordant relative schemes: (i) remove all individuals who have a relative in the data set with the same case-control status, (ii) a greedy matched discordant search in which each case is matched with their most related (not yet matched) control and the case-control pair is selected if the relatedness exceeds 0.20 (otherwise the case is discarded, along with any remaining unmatched controls), and (iii) remove individuals in (i) and then perform the greedy matched discordant search as in (ii). The value of *h*^2^ supplied to LT-Fam for the downsampled data sets was set to 0.257, the H-E regression estimate from the full sample using the thresholded GRM (after transformation to liability scale). We also ran LT-Fam with mis-specified *h*^2^ values ranging from 0.25 and 0.75. All analyses assumed a disease prevalence of 8%, corresponding to a liability threshold of 1.405.

## Results

### Simulated Genotypes and Simulated Phenotypes

We first conducted simulations of sibling pairs using simulated genotypes and simulated case-control phenotypes at different values of disease prevalence under three ascertainment schemes: unbiased, concordant siblings and discordant siblings (see Materials and Methods). We compared the performance of ATT, MLM, LTMLM and LT-Fam. We chose these statistics because a previous study reported that MLM performs at least as well as other methods in family-based association studies^8^ (although that study did not consider family-biased ascertainment). Results are displayed in Table 1 and Table S1. In the unbiased simulations, all of the statistics were roughly correctly calibrated, having mean close to 1 for null SNP sets. In the concordant sibling simulations, LT-Fam was consistently well-calibrated (average *χ*^2^ = 1.00), while the other methods were mis-calibrated. For example, for *N*=1000 and disease prevalence 1%, MLM was severely deflated (average *χ*^2^ = 0.67), and both ATT (1.50) and LTMLM (1.48) were severely inflated. In addition, for *N*=1000 and disease prevalence 1%, LT-Fam attained 3% higher average *χ*^2^ at causal markers vs. MLM and 8% higher average *χ*^2^ at causal markers vs. ATT after calibrating by dividing by the respective *λ*_GC_. In the discordant sibling simulations, LT-Fam was again consistently well-calibrated (average *χ*^2^ = 1.00), while the other methods were again mis-calibrated. For example, for *N*=1000 and disease prevalence 1%, ATT and MLM were severely deflated (average *χ*^2^ = 0.50); LTMLM was only somewhat deflated (0.89) but did not run to completion at other parameter settings. In discordant sibling pair simulations LT-Fam has similar power to other methods after calibrating by dividing by the respective *λ*_GC_. We obtained similar results in simulations with shared environment, based on *ρ*=0.75 between the environmental components of liability of two siblings (Table 2). In the concordant sibling pair simulations, LT-Fam was well-calibrated while MLM was severely deflated and ATT and LTMLM were severely inflated, and LT-Fam attained higher average *χ*^2^ at causal markers after calibrating by dividing by the respective *λ*_GC_. In the discordant sibling pair simulations, LT-Fam was well-calibrated while ATT and MLM were severely deflated and LTMLM did not run to completion, and LT-Fam attained similar average *χ*^2^ at causal markers after calibrating by dividing by the respective *λ*_GC_.

**Table 1.** Simulations under different family-biased ascertainment schemes. We report average association statistics (s.e. in parentheses) at causal markers, null markers and all markers, as well as *λ*_GC_ computed using all markers, for each method at various parameter settings. NA indicates either lack of convergence or a nonsingular phenotypic covariance matrix.

| M | N | Sibling Set | Prevalence | Set | ATT | MLM | LTMLM | LT-Fam |
| --- | --- | --- | --- | --- | --- | --- | --- | --- |
| 50000 | 1000 | Unbiased | 50% | Causal | 16.091(0.668) | 15.423(0.641) | 14.26(0.591) | 14.502(0.601) |
|  |  |  |  | Null | 1.105(0.001) | 1.044(0.001) | 0.965(0.001) | 1.004(0.001) |
|  |  |  |  | All | 1.108(0.001) | 1.047(0.001) | 0.968(0.001) | 1.007(0.001) |
| | | | | All $\lambda_{GC}$ | 1.108(0.002) | 1.047(0.002) | 0.967(0.005) | 1.007(0.002) |
|  |  |  | 10% | Causal | 21.763(0.892) | 20.484(0.839) | 21.67(0.897) | 20.889(0.865) |
|  |  |  |  | Null | 1.062(0.001) | 0.986(0.001) | 1.026(0.001) | 1.004(0.001) |
|  |  |  |  | All | 1.066(0.001) | 0.990(0.001) | 1.030(0.001) | 1.008(0.001) |
| | | | | All $\lambda_{GC}$ | 1.062(0.004) | 0.987(0.004) | 1.026(0.002) | 1.003(0.002) |
|  |  |  | 1% | Causal | 36.139(1.419) | 31.663(1.237) | 39.148(1.576) | 35.62(1.419) |
|  |  |  |  | Null | 1.054(0.001) | 0.921(0.001) | 1.085(0.001) | 1.004(0.001) |
|  |  |  |  | All | 1.061(0.001) | 0.927(0.001) | 1.093(0.001) | 1.011(0.001) |
| | | | | All $\lambda_{GC}$ | 1.055(0.010) | 0.921(0.007) | 1.086(0.002) | 1.005(0.001) |
|  |  | Concord | 50% | Causal | 26.58(1.098) | 12.13(0.512) | 27.751(1.246) | 17.785(0.746) |
|  |  |  |  | Null | 1.502(0.001) | 0.680(0.001) | 1.489(0.001) | 1.001(0.001) |
|  |  |  |  | All | 1.507(0.001) | 0.683(0.001) | 1.494(0.001) | 1.004(0.001) |
| | | | | All $\lambda_{GC}$ | 1.501(0.002) | 0.680(0.001) | 1.489(0.001) | 0.987(0.005) |
|  |  |  | 10% | Causal | 36.048(1.49) | 16.675(0.700) | 27.04(1.193) | 24.767(1.057) |
|  |  |  |  | Null | 1.499(0.001) | 0.676(0.001) | 1.446(0.001) | 0.999(0.001) |
|  |  |  |  | All | 1.506(0.001) | 0.679(0.001) | 1.451(0.001) | 1.003(0.001) |
| | | | | All $\lambda_{GC}$ | 1.499(0.002) | 0.676(0.002) | 1.446(0.004) | 0.986(0.005) |
|  |  |  | 1% | Causal | 67.318(2.393) | 31.717(1.155) | 63.901(2.418) | 47.750(1.773) |
|  |  |  |  | Null | 1.499(0.001) | 0.674(0.001) | 1.475(0.001) | 0.999(0.001) |
|  |  |  |  | All | 1.513(0.001) | 0.680(0.001) | 1.487(0.001) | 1.008(0.001) |
| | | | | All $\lambda_{GC}$ | 1.502(0.002) | 0.674(0.001) | 1.484(0.003) | 0.985(0.005) |
|  |  | Discord | 50% | Causal | 6.354(0.270) | 6.354(0.270) | NA(NA) | 12.272(0.514) |
|  |  |  |  | Null | 0.504(0.001) | 0.504(0.001) | NA(NA) | 1.003(0.001) |
|  |  |  |  | All | 0.505(0.001) | 0.505(0.001) | NA(NA) | 1.005(0.001) |
| | | | | All $\lambda_{GC}$ | 0.516(0.001) | 0.516(0.001) | NA(NA) | 1.059(0.001) |
|  |  |  | 10% | Causal | 9.096(0.376) | 9.096(0.376) | NA(NA) | 17.871(0.736) |
|  |  |  |  | Null | 0.504(0.001) | 0.504(0.001) | NA(NA) | 1.005(0.001) |
|  |  |  |  | All | 0.506(0.001) | 0.506(0.001) | NA(NA) | 1.009(0.001) |
| | | | | All $\lambda_{GC}$ | 0.516(0.001) | 0.516(0.001) | NA(NA) | 1.061(0.002) |
|  |  |  | 1% | Causal | 12.594(0.543) | 12.594(0.543) | 7.057(0.315) | 25.24(1.101) |
|  |  |  |  | Null | 0.503(0.001) | 0.503(0.001) | 0.893(0.001) | 1.004(0.001) |
|  |  |  |  | All | 0.506(0.001) | 0.506(0.001) | 0.895(0.001) | 1.009(0.001) |
| | | | | All $\lambda_{GC}$ | 0.515(0.001) | 0.515(0.001) | 0.893(0.008) | 1.057(0.003) |

**Table 2:** Simulations with shared environment under different family-biased ascertainment schemes. We report average association statistics (s.e. in parentheses) at causal markers, null markers and all markers, as well as *λ*_GC_ computed using all markers, for each method at various parameter settings. NA indicates either lack of convergence or a nonsingular phenotypic covariance matrix.

| M | N | Prevalence | Sibling Set | Set | ATT | MLM | LTMLM | LT-Fam |
| --- | --- | --- | --- | --- | --- | --- | --- | --- |
| 10000 | 1000 | 1% | Unbiased | Causal | 11.855(0.560) | 10.258(0.470) | 6.973(0.273) | 9.749(0.435) |
|  |  |  |  | Null | 1.179(0.002) | 0.987(0.001) | 0.771(0.001) | 1.004(0.001) |
|  |  |  |  | All | 1.190(0.002) | 0.996(0.002) | 0.777(0.001) | 1.013(0.002) |
| | | | | All $\lambda_{GC}$ | 1.187(0.022) | 0.992(0.014) | 0.770(0.006) | 1.010(0.003) |
|  |  |  | Concord | Causal | 43.073(1.703) | 21.365(0.864) | 39.269(1.64) | 29.925(1.238) |
|  |  |  |  | Null | 1.502(0.002) | 0.773(0.001) | 1.393(0.002) | 1.002(0.001) |
|  |  |  |  | All | 1.544(0.003) | 0.794(0.002) | 1.430(0.003) | 1.031(0.002) |
| | | | | All $\lambda_{GC}$ | 1.506(0.004) | 0.774(0.003) | 1.391(0.011) | 1.009(0.005) |
|  |  |  | Discord | Causal | 18.059(0.755) | 18.06(0.755) | NA(NA) | 36.195(1.533) |
|  |  |  |  | Null | 0.504(0.001) | 0.504(0.001) | NA(NA) | 1.005(0.001) |
|  |  |  |  | All | 0.522(0.001) | 0.522(0.001) | NA(NA) | 1.041(0.002) |
| | | | | All $\lambda_{GC}$ | 0.513(0.002) | 0.513(0.002) | NA(NA) | 1.041(0.006) |

LT-Fam results are based on knowledge of the correct *h*^2^ (and did not use REML or H-E regression estimates), whereas other methods are not designed to use this knowledge. We determined that *h*^2^ estimates from both REML and H-E regression were in fact biased in settings of family-biased ascertainment (Table S2 and Table S3), which partially explains the mis-calibration of MLM and LTMLM statistics (Table 1 and Table S1). ^14^^;^ ^15^ Specifically, *h*^2^ was generally overestimated in the concordant sibling simulations, and incorrectly estimated to have value 0 in the discordant sibling simulations (which causes the MLM statistic to become identical to ATT).

Although LT-Fam relies on knowledge of the correct *h*^2^, it is possible that estimates of *h*^2^ obtained from the literature could be incorrect. We thus evaluated the impact of mis-specification of *h*^2^, where the value of 0.25, 0.40, 0.60, and 0.75 supplied to LT-Fam differed from the true value of 0.50, based on concordant sibling pair simulations at a disease prevalence of 1% (where improper calibration was observed for other statistics). We determined that mis-specification of *h*^2^ had virtually no effect either on the calibration of LT-Fam or on the relative value of its average *χ*^2^ at causal markers vs. other methods, after calibrating by dividing by the respective *λ*_GC_ (Table S4).

### CARe Genotypes and T2D Phenotypes

We analyzed 7,088 individuals (1,269 type 2 diabetes cases and 5,819 controls) from the African-American CARe cohort genotyped on genome-wide arrays (see Materials and Methods). We analyzed the full data set and 6 downsampled data sets with family-biased ascertainment: 3 with concordant relatives and 3 with discordant relatives (see Materials and Methods). Results for ATT, MLM, LTMLM, and LT-Fam are displayed in Table 3. In the full data set, LT-Fam, MLM and LTMLM were close to correctly calibrated (we note that average *χ*^2^ slightly larger than 1 may be due to true causal effects^19^) whereas ATT was slightly inflated, as expected due to the family structure in this data. In the concordant relative data sets, LT-Fam was close to correctly calibrated while MLM was deflated (e.g. average *χ*^2^ = 0.82) and ATT was severely inflated. In the discordant relative data sets, LT-Fam was again close to correctly calibrated while ATT and MLM were severely deflated (e.g. average *χ*^2^ = 0.60). These results are similar to what we observed in our simulations (Table 1 and Table S1).

**Table 3.** Results on downsampled CARe T2D samples. We report average association statistics (s.e. in parentheses) at causal markers, null markers and all markers, as well as *λ*_GC_ computed using all markers, for each method. We analyze the entire data set, 3 concordant relative downsampled data sets, and 3 discordant relative downsampled data sets. NA indicates either lack of convergence or a nonsingular phenotypic covariance matrix.

| Down sampling | Sample | N | Number Cases | Number Controls | ATT | MLM | LTMLM | LT-Fam |
| --- | --- | --- | --- | --- | --- | --- | --- | --- |
| None | All | 7088 | 1269 | 5819 | 1.149(0.002) | 1.000(0.002) | 1.001(0.002) | 1.096(0.002) |
| Concord | Removed Cases | 6297 | 478 | 5819 | 1.451(0.002) | 0.983(0.002) | NA(NA) | 1.038(0.002) |
|  | Removed Controls | 5091 | 1269 | 3822 | 1.581(0.003) | 0.960(0.002) | NA(NA) | 1.055(0.002) |
|  | Removed Both | 4300 | 478 | 3822 | 2.534(0.004) | 0.823(0.001) | NA(NA) | 1.074(0.002) |
| Discord | Removed Concord | 2952 | 814 | 2138 | 1.042(0.002) | 0.999(0.007) | 0.979(0.002) | 1.056(0.002) |
|  | Matched Discord | 734 | 367 | 367 | 0.602(0.001) | 0.602(0.005) | NA(NA) | 0.988(0.002) |
|  | Removed Concord and Matched Discord | 176 | 88 | 88 | 0.599(0.001) | 0.599(0.001) | NA(NA) | 1.007(0.002) |

We determined that *h*^2^ estimates from both REML and H-E regression were biased in the downsampled data sets (Table S5), which explains the mis-calibration of MLM statistics in Table 3. Specifically, *h*^2^ was overestimated in the concordant relative data sets, and incorrectly estimated to have value 0 in the discordant relative data sets (which causes the MLM statistic to become identical to ATT), just as in our simulations (Table S2).

## Discussion

We have introduced LT-Fam, a liability threshold mixed model association statistic for family-based case-control studies. In analyses of both simulated concordant/discordant sibling studies and real CARe T2D samples, we have demonstrated that existing association statistics are mis-calibrated under family-biased ascertainment, and that LT-Fam is properly calibrated and attains higher power in some settings.

Initial work on association statistics for family-based case-control studies includes MQLS^20^ and ROADTRIPS.^21^ A recent study determined that standard mixed model association methods^1^^;^ ^2^ perform at least as well in most settings, however, a key advantage of MQLS and ROADTRIPS is that they take advantage of all phenotype information, even for individuals that have not been genotyped. More recently, LTMLM,^5^ LEAP,^6^ and CARAT^12^ (which employ similar ideas) have been developed to address the challenges caused by case-control ascertainment in studies of unrelated individuals. However, to our knowledge, LT-Fam is the first method that addresses the challenges caused by family-biased case-control ascertainment.

Despite its effective modeling of family-biased ascertainment, LT-Fam has several limitations. First, LT-Fam requires published estimates of *h*^2^ from the literature; however, we demonstrated that the method is robust to mis-specification of this parameter (Table S4). Second, LT-Fam requires running time O(*MN*^2^), which is not as fast as state-of-the-art mixed model association methods developed for unascertained samples^4^. Third, LT-Fam does not currently handle fixed-effect covariates, whose inclusion has been shown to be increase power in other settings.^7^^;^ ^22^ Fourth, similar to LTMLM,^5^ the method relies on the assumption of an underlying normally distributed liability; this is widely believed to be a reasonable assumption^11^^;^ ^23^, but may not always hold. Finally, we have not considered analyses of multiple phenotypes.^24^ We nonetheless anticipate that LT-Fam will be a valuable tool in association studies with family-biased case-control ascertainment.

## Acknowledgements

This research was funded by NIH grant R01 HG006399.

## Web Resources

Liability threshold mixed model association statistic for family-based case-control studies (LT-Fam) is updated in the LTMLM software: https://data.broadinstitute.org/alkesgroup/LTMLM/

## References

1. Kang, H.M., Sul, J.H., Service, S.K., Zaitlen, N.A., Kong, S.Y., Freimer, N.B., Sabatti, C., and Eskin, E. (2010). Variance component model to account for sample structure in genome-wide association studies. Nat Genet 42, 348–354.

2. Zhou, X., and Stephens, M. (2012). Genome-wide efficient mixed-model analysis for association studies. Nature genetics 44, 821–824.

3. Yang, J., Zaitlen, N.A., Goddard, M.E., Visscher, P.M., and Price, A.L. (2014). Advantages and pitfalls in the application of mixed-model association methods. Nat Genet 46, 100–106.

4. Loh, P.-R., Tucker, G., Bulik-Sullivan, B.K., Vilhjalmsson, B.J., Finucane, H.K., Salem, R.M., Chasman, D.I., Ridker, P.M., Neale, B.M., Berger, B., et al. (2015). Efficient Bayesian mixed-model analysis increases association power in large cohorts. Nat Genet 47, 284–290.

5. Hayeck, T.J., Zaitlen, N.A., Loh, P.-R., Vilhjalmsson, B., Pollack, S., Gusev, A., Yang, J., Chen, G.-B., Goddard, M.E., and Visscher, P.M. (2015). Mixed Model with Correction for Case-Control Ascertainment Increases Association Power. The American Journal of Human Genetics 96, 720–730.

6. Weissbrod, O., Lippert, C., Geiger, D., and Heckerman, D. (2015). Accurate liability estimation improves power in ascertained case-control studies. Nature methods 12, 332–334.

7. Jiang, D., Zhong, S., and McPeek, M.S. (2016). Retrospective Binary-Trait Association Test Elucidates Genetic Architecture of Crohn Disease. Am J Hum Genet 98, 243–255.

8. Eu-ahsunthornwattana, J., Miller, E.N., Fakiola, M., Jeronimo, S.M., Blackwell, J.M., Cordell, H.J., and 2, W.T.C.C.C. (2014). Comparison of methods to account for relatedness in genome-wide association studies with family-based data.

9. Zaitlen, N., Kraft, P., Patterson, N., Pasaniuc, B., Bhatia, G., Pollack, S., and Price, A.L. (2013). Using extended genealogy to estimate components of heritability for 23 quantitative and dichotomous traits.

10. Xia, C., Amador, C., Huffman, J., Trochet, H., Campbell, A., Porteous, D., Hastie, N.D., Hayward, C., Vitart, V., Navarro, P., et al. (2016). Pedigree-and SNP-Associated Genetics and Recent Environment are the Major Contributors to Anthropometric and Cardiometabolic Trait Variation. PLoS Genet 12, e1005804.

11. Visscher, P.M., Hill, W.G., and Wray, N.R. (2008). Heritability in the genomics era–concepts and misconceptions. Nat Rev Genet 9, 255–266.

12. Listgarten, J., Lippert, C., Kadie, C.M., Davidson, R.I., Eskin, E., and Heckerman, D. (2012). Improved linear mixed models for genome-wide association studies. Nature methods 9, 525–526.

13. Falconer, D.S. (1967). The inheritance of liability to diseases with variable age of onset, with particular reference to diabetes mellitus. Ann Hum Genet 31, 1–20.

14. Haseman, J., and Elston, R. (1972). The investigation of linkage between a quantitative trait and a marker locus. Behavior genetics 2, 3–19.

15. Golan, D., Lander, E.S., and Rosset, S. (2014). Measuring missing heritability: Inferring the contribution of common variants. Proceedings of the National Academy of Sciences 111, E5272–E5281.

16. Lee, S.H., Wray, N.R., Goddard, M.E., and Visscher, P.M. (2011). Estimating missing heritability for disease from genome-wide association studies. Am J Hum Genet 88, 294–305.

17. Devlin, B., and Roeder, K. (1999). Genomic control for association studies. Biometrics 55, 997–1004.

18. Pasaniuc, B., Zaitlen, N., Lettre, G., Chen, G.K., Tandon, A., Kao, W., Ruczinski, I., Fornage, M., Siscovick, D.S., and Zhu, X. (2011). Enhanced statistical tests for GWAS in admixed populations: assessment using African Americans from CARe and a Breast Cancer Consortium. PLoS Genet 7, e1001371.

19. Yang, J., Weedon, M.N., Purcell, S., Lettre, G., Estrada, K., Willer, C.J., Smith, A.V., Ingelsson, E., O’Connell, J.R., and Mangino, M. (2011). Genomic inflation factors under polygenic inheritance. European Journal of Human Genetics 19, 807–812.

20. Thornton, T., and McPeek, M.S. (2007). Case-control association testing with related individuals: a more powerful quasi-likelihood score test. Am J Hum Genet 81, 321–337.

21. Thornton, T., and McPeek, M.S. (2010). ROADTRIPS: case-control association testing with partially or completely unknown population and pedigree structure. Am J Hum Genet 86, 172–184.

22. Zaitlen, N., Lindström, S., Pasaniuc, B., Cornelis, M., Genovese, G., Pollack, S., Barton, A., Bickeböller, H., Bowden, D.W., and Eyre, S. (2012). Informed conditioning on clinical covariates increases power in case-control association studies. PLoS Genet 8, e1003032.

23. Plomin, R., Haworth, C.M., and Davis, O.S. (2009). Common disorders are quantitative traits. Nature Reviews Genetics 10, 872–878.

24. Zhou, X., and Stephens, M. (2014). Efficient algorithms for multivariate linear mixed models in genome-wide association studies. Nature methods 11, 407.

